# Bacterial swarm-mediated phage transportation disrupts a biofilm inherently protected from phage penetration

**DOI:** 10.1101/2021.06.25.449910

**Authors:** Nichith K. Ratheesh, Cole A. Calderon, Amanda M. Zdimal, Abhishek Shrivastava

## Abstract

The treatment of chronic bacterial infections by phages has shown promise in combating antimicrobial resistance. A typical phage particle is at least an order of magnitude larger than an antibiotic molecule. Hence, phages diffuse slower than antibiotics, and can also get trapped in the polymeric mesh of biofilm matrix. By tracking fluorescently labeled lambda phages that do not infect *Capnocytophaga gingivalis*, a bacterium abundant in the human oral microbiota, we demonstrate active phage transportation by a *C. gingivalis* swarm. As a result, the rate of disruption of the prey of lambda phage i.e., an *Escherichia coli* colony, increases 10 times. *C. gingivalis* drills tunnels within a curli fiber containing *E. coli* biofilm and increase the efficiency of phage penetration. This provides evidence for a novel mechanism of phage-bacterial warfare.

## Introduction

Antimicrobial resistance is a global public health threat^1^ and the development of new antibiotics is a glacially slow process. Formation of biofilm by bacterial cells results in a 10-1000 fold increase in antibiotic resistance^2^. Phages are viruses that kill bacteria and are promising alternatives for antibiotic treatment during chronic antimicrobial resistant infections^3–11^. However phage therapy has several limitations that need to be resolved before it becomes a widely applied medical strategy. One limitation is that phages can get trapped in the matrix of infectious biofilms and this diminishes their frequency of encountering a bacterial cell. Another limitation is that the larger size of phages compared to antibiotic molecules results in a slower diffusion rate. According to the Stokes-Einstein equation, the translation diffusion coefficient (*D*) of a sphere of radius *r* in a medium of viscosity *η* is 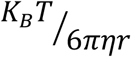 where *K*_*B*_ is the Boltzmann’s constant and *T* is the absolute temperature. *D* is proportional to the speed of diffusion and the inverse relationship between *D* and *r* implies that small molecules diffuse faster. One of the largest known antibiotics, vancomycin (∼1 nm radius), has 66 C atoms^12^. The carbon chain lengths of commonly used medicinal antibiotics are about a quarter to half of vancomycin. For example, amoxicillin and cefalexin have 16 C atoms, ciprofloxacin and methicillin have 17 C atoms, and azithromycin has 38 C atoms^13–16^. The capsid of lambda phage used in our study has a radius of about 30 nm^17^. This suggests that Lambda phage will diffuse 30 times slower than vancomycin, and 100 times slower than cefalexin. Additionally, it has been experimentally demonstrated that phage T4 and fNEL diffuse slower as compared with ciprofloxacin, penicillin, and tetracycline^18^. To establish phage therapy as a widely used efficient alternative to antibiotic treatment, one needs to overcome its diffusion-based limitation.

Theoretically, one way to overcome this limitation is by providing a directionality to diffusion. Electron microscopy images have shown that phages bind to bacterial flagella^19^, and analysis of bacterial colonies suggests hitchhiking of phages^19,20^. However, live microscopy images of phages hitchhiking on bacteria are not available, and direct data for the ‘hiking’ aspect of ‘hitchhiking’ are missing. Can one directly visualize phage transportation? Also, are motile but non-flagellated bacteria able to transport phages? We sought to answer these questions with the help of *Capnocytophaga gingivalis*, which is a member of the bacterial phylum Bacteroidetes. *C. gingivalis* exhibits gliding motility and transports non-motile bacterial species of the human oral microbiota^21^.

Swimming, twitching, and gliding are the three major modes of motility exhibited by bacteria^22,23^. Bacterial gliding is an active process and individual rod-shaped gliding bacteria move in a screw-like fashion^24,25^. Bacteria of the phylum Bacteroidetes are among the abundant members of the healthy human microbiome and motile members of the phylum Bacteroidetes perform gliding motility with the help of the Type 9 Secretion System (T9SS) which is a rotary motor that couples with a cell-surface track^26–28^. SprB is loaded on the track and its interaction with an external substratum results in gliding motility^29^. On an agar surface, T9SS driven cells of *C. gingivalis* and other gliding Bacteroidetes swarm in a vortex-like fashion^21,30^. Particle velocimetry of tracer gas bubbles has shown that the swarm fluid moves along the direction of a *C. gingivalis* swarm^21^.

## Results

Our data show that a swarm of *C. gingivalis*, a bacterium that is abundant in the oral microbiota of healthy humans^31^, can actively transport lambda phages over long distances (**Movie 1** and **Fig 1a, b**). The experiments were performed using lambda phages that have Yellow Fluorescent protein (YFP) attached to their coat protein gpD^32^. Lambda phage infects *E. coli* but cannot infect *C. gingivalis* (**Fig S1**). We found that a swarm of *C. gingivalis* can be used as a tool for phage delivery.

**Fig 1.**
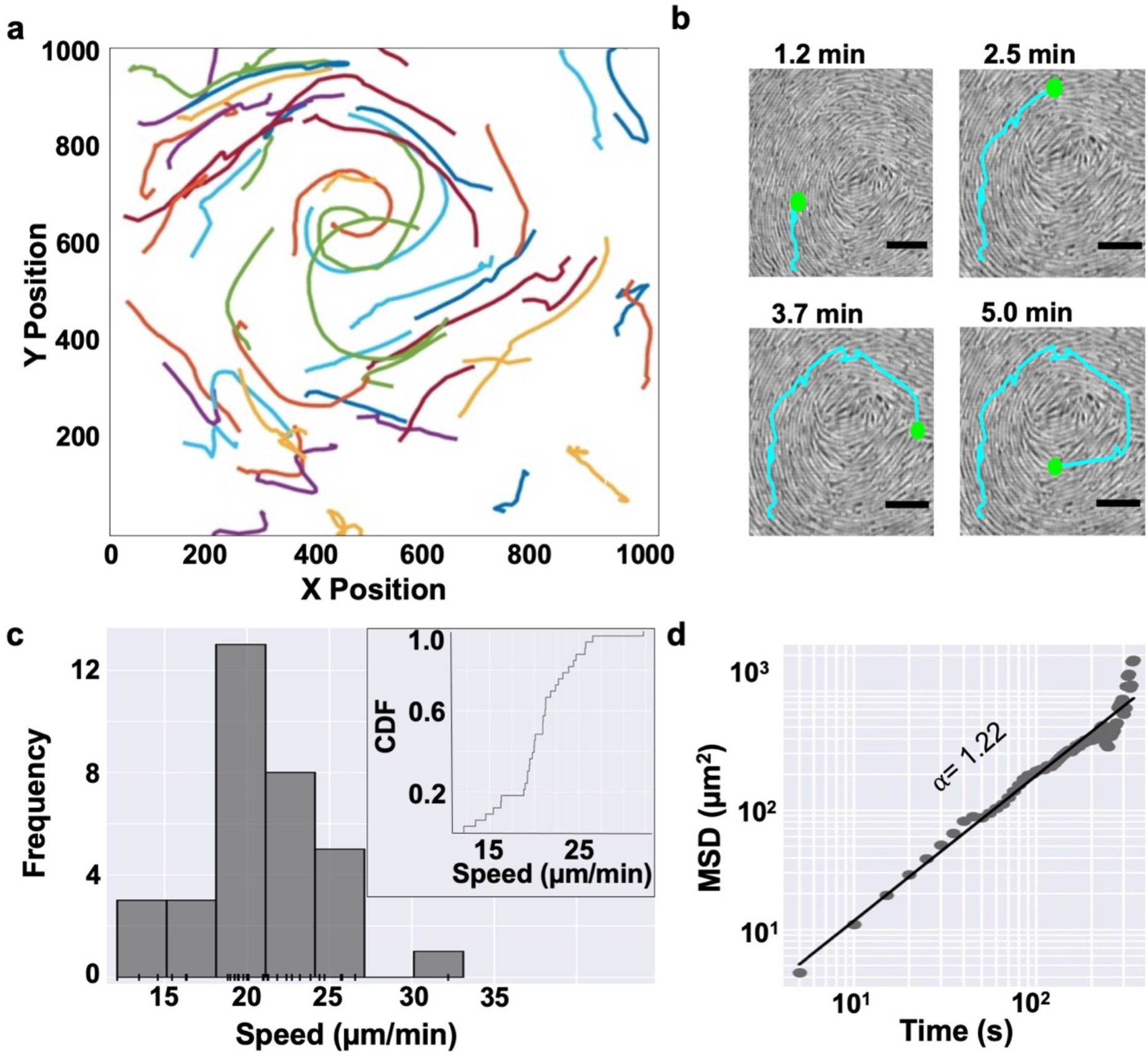
A swarm of *C. gingivalis* actively transports phages. **a**, Trajectories from **Movie 1** show 33 phages being propelled by a *C. gingivalis* swarm. **b**, Time-lapse images of a cropped section of **Movie 1** showing fluorescent lambda phage being transported by a swarm of *C. gingivalis* (grey). Position of a phage and the trajectory are shown in green and cyan, respectively. Scale bar = 5 μm. **c**, A frequency distribution and a rug plot of phage speed. Inset shows the cumulative density function (CDF) of phage speed. **d, Ensemble** Mean squared displacement of phages from **Movie 1** plotted as a function of time. The slope (α = 1.22) of the power-law fit implies that the phages are super diffusive and are actively propelled by a *C. gingivalis* swarm.

Phages were transported due to hydrodynamic fluid flows generated by *C. gingivalis* swarm (**Movie 1**). In most cases, a phage did not bind to a *C. gingivalis* cell (**Movie 2**). In very rare cases, transient attachment of a phage to a *C. gingivalis* cell was observed (**Movie 3** and **Fig S2**). A bacterial swarm secretes osmotic agents that form a gradient due to which water moves out from the agar, mixes with surfactants, and forms the swarm fluid^33^. Surfactants secreted into the swarm fluid keep the swarming cells wet and help in reduction of water loss from the substratum^34^. By using microbubbles as tracers, it was shown that *E. coli* swarm fluid moves orthogonal to the swarm front’s direction of motion^33^. This was attributed to the direction of rotation of the *E. coli* flagella bundle being orthogonal to the long axis of the cell. In the case of gliding bacterium, the mobile adhesin, SprB, moves along the long axis of the cell. Hence, microbubbles move along the direction of motion of a *C. gingivalis* swarm^21^. Tracking showed that phages were transported by *C. gingivalis* in the direction of the swarm’s propagation (**Fig 1a, b**) with a mean speed of 20.71 +/- 3.96 μm/minute (**Fig 1c**), which is also the speed of motion of individual cells in a swarm^21^. Changes in Mean Square Displacement (MSD) as a function of time provided a power-law slope of 1.22 **(Fig 1d)**. MSD of 98 phages from the 12 additional swarms (**Fig S3**) was similar to the value reported in **Fig 1d**. Hence, the transport of phages by a *C. gingivalis* swarm is an active process.

To test if swarm-mediated phage transport provides an advantage over phage transport by diffusion, we performed simulations that are based on Fick’s laws of diffusion. The predictions of our simulations were tested experimentally. To design a control for our experiment, it was necessary to find the diffusion constant of phages within a thin liquid layer on the wet agar surface. We inoculated 2 μL of 10^8^ PFU/mL fluorescent lambda phages over an agar surface that was incubated in a 100% relative humidity chamber. Diffusing fluorescent phages in a thin liquid on the agar surface were imaged (**Movie 4**) and their MSD power-law slope was 0.68 (**Fig 2a**), which suggests that the phages were sub-diffusive. Using the Einstein relation^25^ 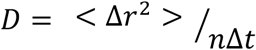 we calculated the diffusion coefficient (*D*) of phages on a wet agar surface to be 0.041 μm^2^s^-1^. Similarly, we calculated that *D* of phages on swarm fluid of bacteria with inhibited motility is 0.228 (**Fig 2a**). Here, Δ*r*^2^ is the change in squared displacement, *n* is the dimensionality of the diffusion, and Δ*t* is the change in time^35^. To measure the distance that phages will diffuse in 30 hours at 33°C, we used Fick’s second law of diffusion^35^ 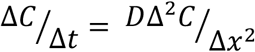 where Δ*C* is the change in concentration, Δ*t* is the change in time, *D* is the diffusion coefficient and Δ*x* is the change in distance. We found that within 30 hours, phages diffuse about 400 μm on a wet agar surface. In contrast, they diffuse only 50 μm on the swarm fluid of *C. gingivalis* with inhibited motility. (**Fig 2b**). *C. gingivalis* swarms in a circular fashion wherein new circles propagate from the edge of an initial circular swarm^21^. The vortexing speed of the swarm is about 20 μm/min. Using equations 1-3 shown in **Fig 2c**, the linear speed was calculated to be 6.36 μm/min. Using the advection-diffusion equation^35^ 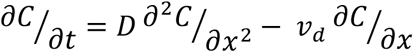 we simulated that *C. gingivalis* can transport phages to a distance of ∼1500 μm in 3.3 hours (**Fig 2d**). Here, *c* is the concentration of phages, *t* is time, *x* is distance, *D* is the diffusion coefficient, and *v*_*d*_ is advective velocity.

**Fig 2.**
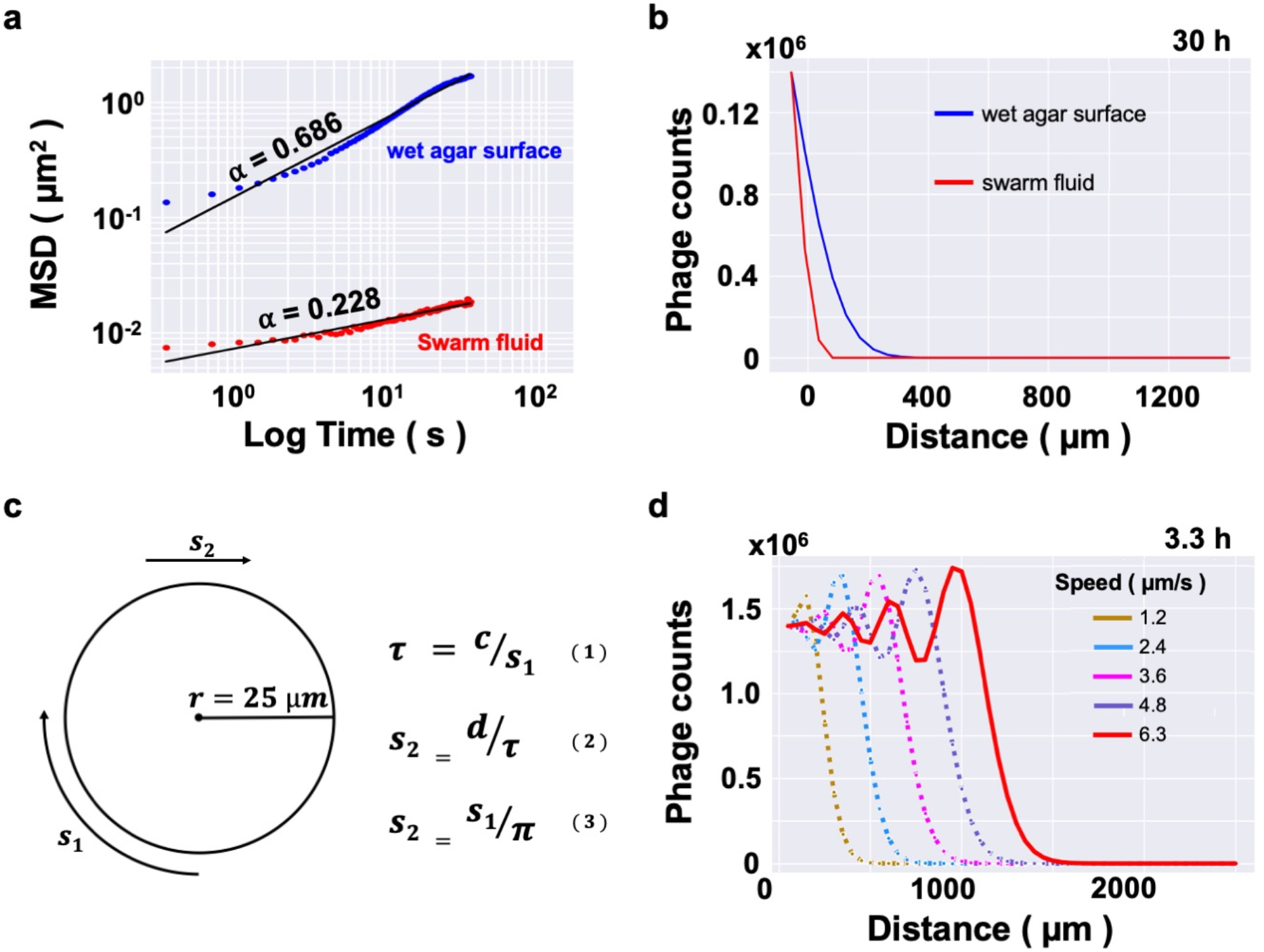
Prediction of the distance travelled by transported phages. **a**, Mean squared displacement of phages diffusing in a thin layer of liquid on a wet agar surface (blue) and swarm fluid of bacteria with inhibited motility (red). **b**, A prediction of one-dimensional distance covered by phages diffusing in the two fluids described above. **c**, A cartoon outlining a vortexing swarm with radius *r*, circumference *c*, and vortexing speed *s*_1_. Using equations 1-3, linear speed s_2_ is calculate**d** to be 6.3 μm/min. **d**, A prediction of one-dimensional distance covered by actively transported phages. Outputs for linear speed ranging from 1.2 μm/min to 6.3 μm/min are shown.

The in-silico results described in **Fig 2** guided the design of the wet-lab experiments described in **Fig 3**. We know that *C. gingivalis* starts swarming about 7 hours after initial inoculation on an agar surface. Hence, we started capturing microscopic images to test if phages delivered by *C. gingivalis* can increase the kinetics of *E. coli* colony disruption 10 hours after the initial inoculation. As suggested by the simulation in **Fig 2d**, this provided 3 hours for phage delivery and 7 hours for swarm initiation.

**Fig 3.**
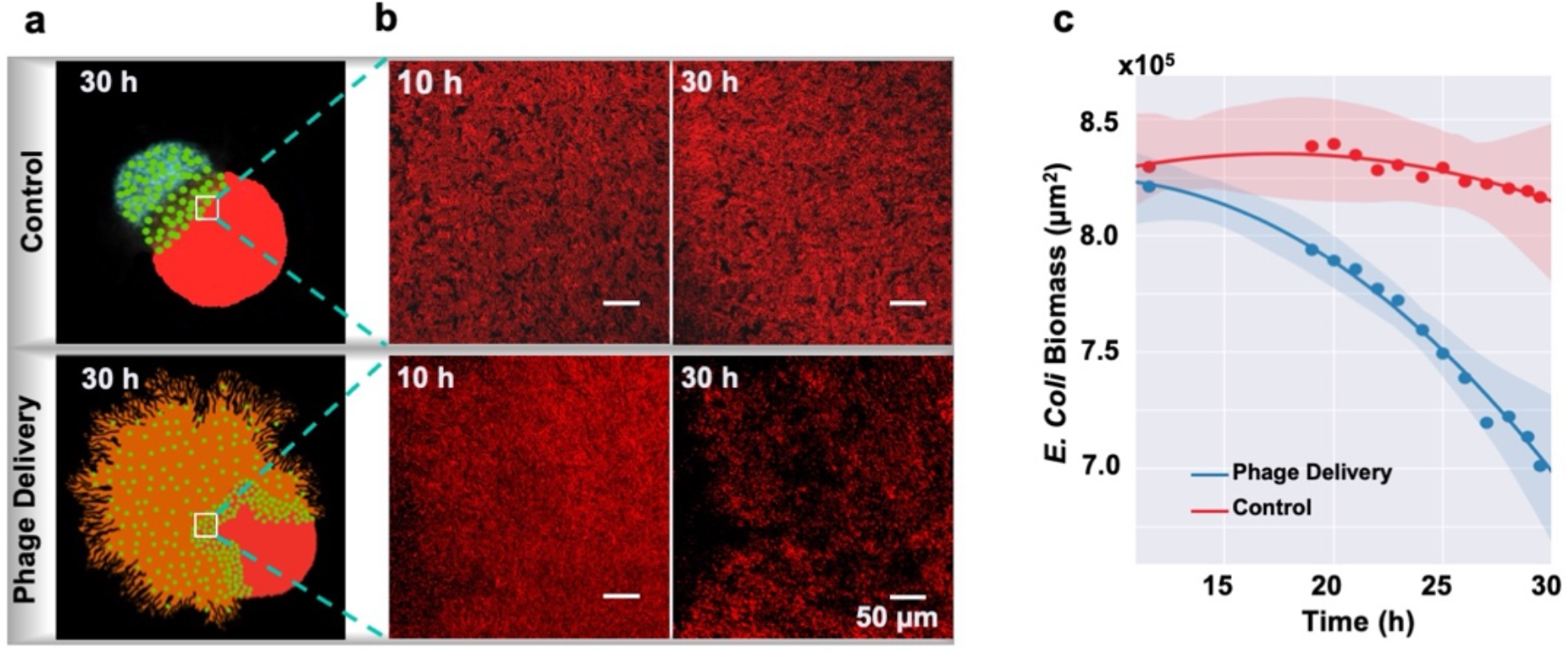
Phage delivery reduces biomass of the prey. **a**, Cartoons depicting control and experimental settings. Phages, *E. coli*, and *C. gingivalis are in green, red, and orange, respectively*. **b**, Images of the red fluorescent *E. coli* colony shown at two time points for the control and experimental settings. These images accompany **Movies 6-8** and **Fig S4 – S7**. **c**, Changes in the area of *E. coli* biomass depicted as a function of time. Dots represent the mean of three biological replicates. Dark lines are second order regression fits and the light shaded regions represent the 95% confidence interval.

We inoculated phage-CG mix (see Methods) and red fluorescent non-motile *E. coli* (*ΔfliC* with mCherry) on a wet agar surface at 1000 μm from one another. Via confocal microscopy, changes in *E. coli* biomass were measured between 10 to 30 hours. As controls, similar amounts of either lambda phage or *C. gingivalis* suspension were spotted at 1000 μm from the fluorescent *E. coli* colony. Samples were imaged 500 μm inwards from the edge of the *E. coli* colony. In this region, minimal reduction in *E. coli* biomass was observed in the control (**Fig 3a, b, Movie 6**). This implies that artificial fluid flows that might occur during sample preparation and addition of cover glass cannot deliver phages to this region. In contrast, when phages were actively transported by *C. gingivalis*, the *E. coli* biomass reduction occurs 10 times faster (**Fig 3a, b, c and Movie 7**). No change in *E. coli* biomass was observed when *C. gingivalis* alone was spotted 1000 μm from the *E. coli* colony (**Fig S6 - S7** and **Movie 8**). This control shows that inter-bacterial competition due to metabolites or physical forces do not play a role in the reduction of biomass of *E. coli*. The phenotype that we report serves as a novel example of predator-prey relationship in the microbial world.

Recently, Vidakovic and co-workers demonstrated that a polymeric mesh generated by curli fiber creates a physical barrier that protects a biofilm from phage infection^26^. In order to test if active phage transportation can bypass the physical barrier created by curli fiber, we designed a control where we inoculated phages on top of a curli fiber producing *E. coli* biofilm and found that most phages were concentrated at the top layer of the biofilm (**Fig 4a-c**). In contrast, effective phage penetration was observed when phage-CG mix was inoculated at the circumference of a curli fiber-producing mature *E. coli* biofilm. Strikingly, actively transported phages even reached the bottom surface of the biofilm (**Fig 4d-f**). The largest number of phages were found at a depth of about 50 μm of a biofilm that has a total height of 105 μm (**Fig 4i**).

**Fig 4.**
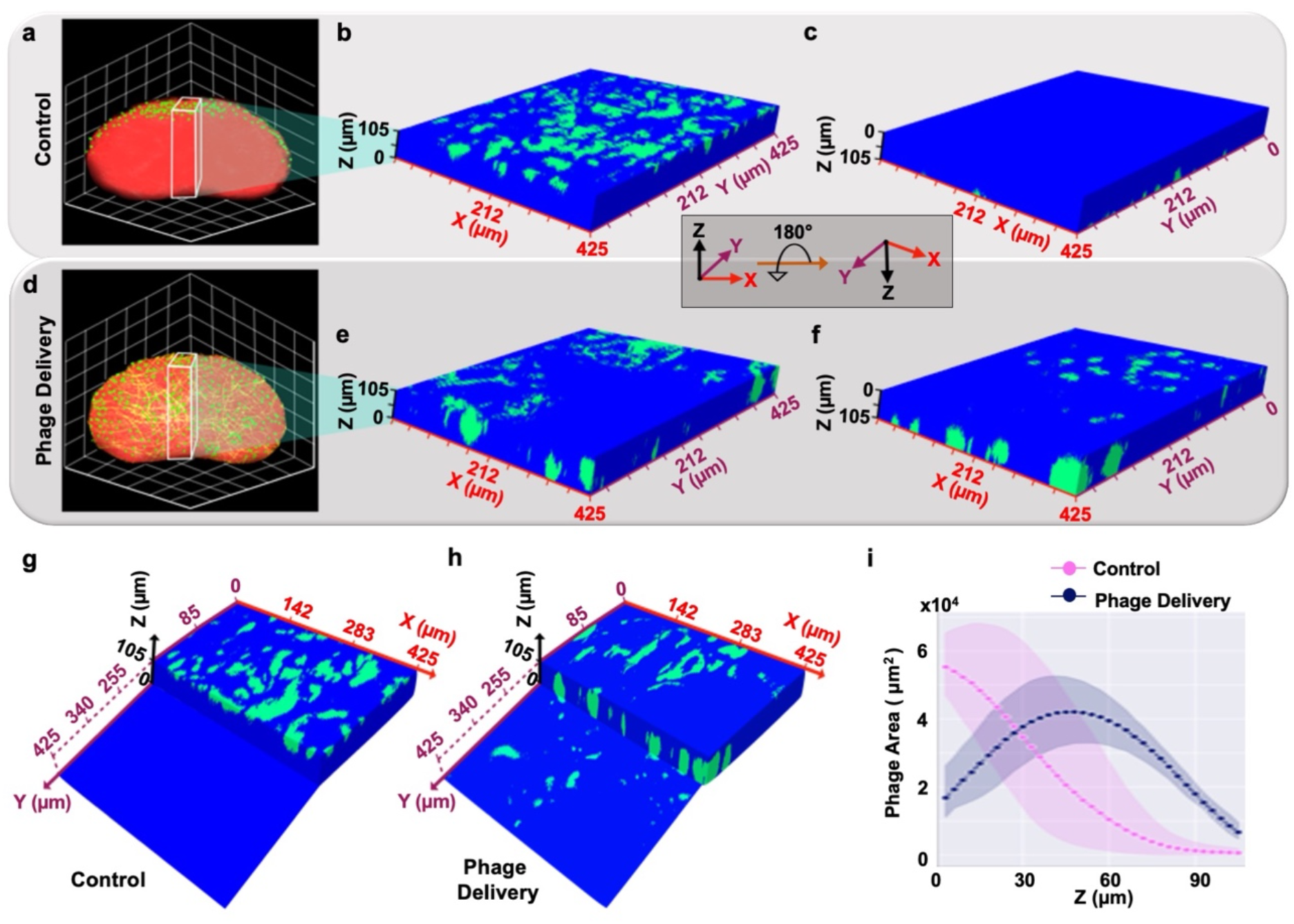
Transport by *C. gingivalis* increases three-dimensional phage delivery within a mature *E. coli* biofilm. **a**, A cartoon showing a curli fiber containing 72-hour old *E. coli* biofilm (red) with phages (green). **b**, Top view and **c**, bottom view of the control. Phages are in green, and blue depicts the rest of the biofilm. These images are taken from **Movie 9. d**, A cartoon of a curli fiber containing 72 h old *E. coli* biofilm (red) with phages (green) being delivered by *C. gingivalis* (yellow). **e**, Top view and **f**, bottom view of the biofilm with phages in green and blue depicts the rest of the biofilm. These images are taken from **Movie 10. g**, A slice of the biofilm allowing partial view of the top and bottom layer for the control and **h**, the experimental setting. **i**, Change in the area of phages along the z-axis show that after delivery by *C. gingivalis*, phages are found in all regions with the maximum numbers being present at a depth of ∼50 μm. Connected dots represent the mean from three biological replicates. The light shaded regions represent the 95% confidence interval. In the control, the greatest density of phage particles is found in the top layer (z = 0 μm).

*C. gingivalis* cells were introduced along the circumference of a fluorescent curli fiber-producing *E. coli* biofilm. After 24 hours of swarming, *C. gingivalis* cells were stained by fluorescence *in situ* hybridization (FISH). Analysis of the imaged data showed that *C. gingivalis* formed tunnels within a 50 μm thick zone of the *E. coli* biofilm (**Fig 5a, b**). The total height of the *E. coli* biofilm was 105 μm and the zone preferred by *C. gingivalis* spans from 20 μm depth to 70 μm depth from the top of the *E. coli* biofilm. This is similar to the depth reported for maximum phage penetration in **Fig 4i**. Single cells of *C. gingivalis* and other closely related bacteria move in a screw-like fashion^21,24^. Our results point towards a mechanism for phage delivery where a self-propelled screw drills tunnels in a biofilm and drives a thin stream of fluid that carries phages to their targets (**Fig 5c, d**). A bacterial swarm contains multiple self-propelled screws, and our data suggests that they might cooperate and work towards drilling a common tunnel.

**Fig 5.**
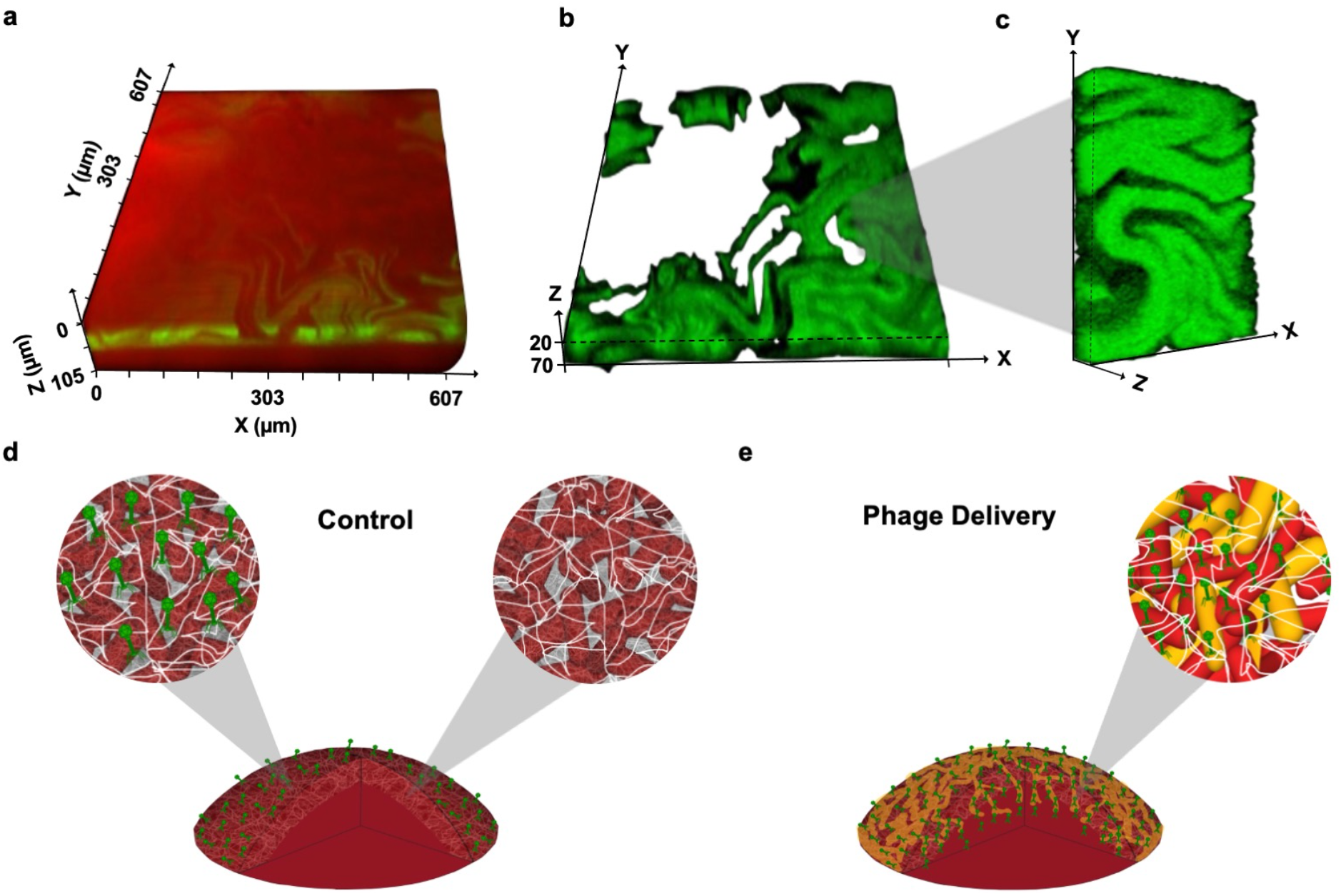
*C. gingivalis* drills tunnels within a mature *E. coli* biofilm. **a**, A three-dimensional view of the invasion of a mature *E. coli* biofilm by *C. gingivalis. E. coli* is shown in red and *C. gingivalis* is shown in green. This image is taken from **Movie 11. b**, An image showing only the *C. gingivalis* channel from panel a. This image is taken from **Movie 12. c**, A zoomed in view of a small section from the image in panel b depicts finger like projections formed by the *C. gingivalis* swarm within a mature *E. coli* biofilm. **d**, A cartoon of the control where due to curli fiber (white), phages (green) fail to penetrate the *E. coli* biofilm (red). **e**, A cartoon depicting the experimental setting where phages (green) can penetrate the biofilm through tunnels formed by the *C. gingivalis* swarm (yellow).

## Discussion

We report that active transportation of phages by a bacterial swarm increases the frequency of interaction between the predator and the prey. This results in an increase in the clearance of biomass of an *E. coli* colony. The transporter *C. gingivalis* forms tunnels within an *E. coli* biofilm and delivers phages to previously inaccessible regions of the biofilm.

While our work improves the fundamental understanding of microbial ecology, in future, it can also be applied to improve the pharmacokinetics of phage therapy. Topical application of phage is the preferred method for treating burn wound infection^36^ and chronic ear infection (otitis media)^8^. Our results demonstrate improved delivery of topically applied phages within an in-vitro *E. coli* biofilm. In future, development of a universal phage transporter might provide an approach for efficient delivery of all phages in the cocktail. Furthermore, chemotactic control of the directionality of a swarm can also help expand active phage delivery to chronic biofilms that grow in relatively inaccessible locations within the human body.

*C. gingivalis* is an understudied bacterium and very little is known about its chemotaxis pathway. The genome of *C. gingivalis* encodes a homolog of the chemoreceptor Aer^37^ and *C. gingivalis* swarms best in CO_2_ rich microaerophilic conditions. It might be possible that via Aer, *C. gingivalis* senses intracellular energy levels and is guided towards a location that allows for a preferred redox environment. Chemotaxis of flagellated bacteria is well-studied and is known to impact autoaggregation^38,39^, symmetrical pattern formation and traveling wave formation^40–42^. Increased physical interactions between cells in a swarm reduces the ability for tracking chemotactic gradients^43^ but the chemotaxis pathway can also be remodeled for swarming^44^. We demonstrate that most of the tunneling *C. gingivalis* are found between the depth of 20 to 70 μm of the *E. coli* biofilm. It might be possible that chemotaxis of the transporter drives preferential localization of the predator within a biofilm of prey.

Swarming *C. gingivalis* generate an active flow within their swarm fluid^21^ and we find that phages can travel along this flow. In an alternate scenario, if *C. gingivalis* could bind to specific phage proteins, the steric hindrances might impede screw-like motility of the transporter cell. Thereby, the overall efficiency of transport might decrease. Thus, it appears that *C. gingivalis* have evolved an efficient transport mechanism. Since phages do not need to bind to *C. gingivalis*, there is a possibility of transport of diverse phages via this mechanism. The natural habitat of *C. gingivalis* is the human oral microbiota and it does not escape our attention that bacterial swarms on gut and oral mucosal surfaces within the human body might be actively transporting virus particles that impact human health.

## Methods

### Growth of *C. gingivalis*

Motile *C. gingivalis* ATCC 33624 was grown on TSY-agar (TSB 30 g/L, yeast extract 3 g/L, and 1.5% Difco Bacto agar), as described previously^21^. Growth was carried out in a CO_2_ rich anaerobic environment with 100% relative humidity. Briefly, inoculated agar plates were placed in an AnaeroPack System jar (Mitsubishi Gas Chemical) with candles, an ignited sheet of Kimtech paper, and a beaker full of water. The box was sealed and incubated at 37°C for 48 h. This is the optimum condition for *C. gingivalis* motility. A *C. gingivalis* suspension was prepared by scraping 27 μg of *C. gingivalis* (wet weight) from a plate and resuspending it in 50 μL of sterile and double distilled water.

### Preparation of phage lysate

Fluorescent phage λ_LZ641_ (gift from Lanying Zeng, Texas A&M University) used in this study contains a mixture of wild-type head stabilization protein gpD and a fusion of gpD with yellow fluorescent protein (eYFP)^45^. *E. coli* strain MG1655 was grown in Luria-Bertani (LB) agar and broth with overnight shaking at 37°C. 100 μL of the *E. coli* culture was mixed with 100 μL of either 10^−6^ or 10^−7^ dilution of a 10^8^ PFU/mL lysate of λ_LZ641_. After incubation at room temperature for 15 minutes, the mixture was added to 4 mL top agar (LB with 0.7% agar) (50°C), gently mixed, and poured on LB agar. After overnight incubation at 37°C, plaque containing top agar was resuspended in 1.2 mL of sterile water. After incubation for 4 hours at room temperature the slurry was centrifuged at 1900 rcf for 5 minutes. The supernatant was passed through a 0.2 μm filter and either kept at 4°C for immediate use or at -20°C for long-term storage. For the plaque assay shown in **Fig S1**, a method like the one described above was used with one notable exception – instead of LB, TSY was used for both *E. coli* and *C. gingivalis*.

### *E. coli* growth conditions

*E. coli* was grown on LB agar and broth. The plasmid pBT1 mCherry (a gift from Michael Lynch, Addgene plasmid # 65823) was inserted via electroporation into non-motile *E. coli* (*ΔfliC* in RP437 background). Cells containing pBT1-mCherry were cultured on LB agar with 100 μg/mL ampicillin at 37°C. A single colony was inoculated in 3 mL LB broth with ampicillin at 37°C and was grown overnight with shaking. Subsequently, a day culture was prepared by adding 300 μL of the overnight culture to 30 mL of LB broth with ampicillin and cells were grown to mid-log phase at 37°C while shaking. The day culture was used for subsequent experimentation.

Curli fiber producing *E. coli* WR3110 strain^46^ (gift from Knut Drescher, University of Basel) was grown on LB agar at 37°C. Like the method described above, plasmid pBT1 mCherry was inserted into *E. coli* WR3110. To test curli fiber production by a biofilm of *E. coli* WR3110, it was grown on TSY agar supplemented with an amyloid dye thioflavin S^44^ (40gml^-1^)^47^ for 48 hours at 28°C. After incubation, mature biofilms were imaged using Zeiss LSM 880 confocal microscope (Oberkochen, Germany) and z-stack images (3 μm slices) of the *E. coli* biofilm were obtained. To visualize the distribution of curli fibers within the biofilm. The maximum intensity projection from z-stacks were reconstructed using the Zeiss Zen software^48^.

### Imaging and analysis of phage transportation

10 μl of fluorescent λ_LZ641_ phage lysate was added to 20 μL of a C. *gingivalis* suspension, we refer to this as the phage-CG mix. 2 μL of the phage-CG mix was spotted on a TSY agar pad on a slide and incubated for 5 minutes at room temperature. Subsequently, a coverslip was placed, and the edges were sealed using beeswax (Aqua Solutions, Inc. Lot#: 730701). The slides were incubated at 37°C for 3 hours in conditions optimum for *C. gingivalis* motility (described above).

Images were captured with the Zeiss LSM880 confocal microscope and phages were tracked using an ImageJ plugin TrackMate^49,50^. The two-dimensional position coordinates of each phage particle were exported from ImageJ in a .xml format. The .xml files were imported into MATLAB^51^, and converted to a .csv file. The .csv file was imported into Python and converted to a DataFrame. The mean speed of each phage particle were calculated using a custom MATLAB script and the mean square displacement (MSD) was calculated using a custom Python script based on TrackPy^52^.

To test if phages attach to individual *C. gingivalis* cells, 50 μL phage-CG mix was injected into a tunnel slide. The tunnel slide was incubated at room temperature for 5 minutes and 50 μL of 10% methyl cellulose was added as described previously^29^. Motile *C. gingivalis* cells on a cover glass and fluorescent phages were imaged by a Zeiss LSM880 confocal microscope.

### Measurement of diffusion coefficient of phage particles and simulation of phage delivery

To calculate the diffusion coefficient of phages on a wet agar surface, 2μl of 10^8^ PFU/ml λ_LZ641_ were spotted on a freshly prepared TSY agar pad and incubated for 5 minutes at room temperature. Subsequently, a coverslip was placed, and the edges were sealed using beeswax. Timelapse images of phages diffusing within a thin liquid layer between the agar and coverglass were captured with the Zeiss LSM880 confocal microscope. Using custom Python scripts, ensemble MSD of phage particles was calculated.

In order to calculate the diffusion coefficient of phages diffusing in swarm fluid of bacteria with inhibited motility, a 48-hour old culture of *C. gingivalis* was treated with 10 μM carbonyl cyanide-m-chlorophenylhydrazone (CCCP), an uncoupler which disrupts the proton motive force and stops T9SS driven motility^21,53^. CCCP treated *C. gingivalis* was used to prepare a phage-CG mix which was spotted on a TSY agar pad. As described above, ensemble MSD of phage particles diffusing in swarm fluid of non-motile *C. gingivalis* was calculated.

To find how far apart *E. coli* and phage-CG mix should be spotted on a TSY agar plate, we performed simulations of phages diffusing on wet agar surface and swarm fluid of bacteria with inhibited motility. Additionally, the distance traveled due to active transportation of phages by a swarm of *C. gingivalis* was also simulated.

### Change in biomass of *E. coli* colony after phage delivery

A day culture of non-motile *E. coli (ΔfliC* in RP437) containing plasmid pBT1 mCherry was pelleted and resuspended in 250 μL of sterile water. 2 μL of the suspension was spotted on a petri dish containing TSY-agar. 2 μL of phage-CG mix was spotted at 1000 μm from the *E. coli* spot. The plate was incubated for 10 hours in conditions optimum for *C. gingivalis* motility (described above). After incubation, a circular slab of agar (around 35 mm diameter) surrounding the two colonies was cut with a scalpel and transferred onto the lid of a smaller petri plate (35 × 10 mm). A coverslip was placed, and the edges were sealed using beeswax. Using a Zeiss LSM880 confocal microscope, time-lapse images were obtained for the next 21 hours. The interval between each captured image was 1 hour. The area of *E. coli* colony was calculated using the Python library Scikit-image^54^. Change in biomass of *E. coli* after phage delivery was determined via a custom Python script.

### Assay of phage delivery within a mature *E. coli* biofilm

A curli-fiber producing *E. coli* strain WR3110 with plasmid pBT1 mCherry was grown on TSY-agar petri plate for 48 hours at 28°C. The lower temperature maximizes curli-fiber production and biofilm formation^55^. After incubation, a 50 μL *C. gingivalis* suspension was pipetted around the circumference of the mature *E. coli* biofilm and the plate was transferred for a 14-hour incubation in conditions optimum for *C. gingivalis* motility (described above). After incubation, 20 μL of 10^5^ PFU/mL λ_LZ641_ was spotted on the biofilm and the plate was incubated for another 10 hours in conditions optimum for *C. gingivalis* motility (described above). After incubation, z-stack images (3 μm slices) of the *E. coli* biofilm were obtained using a Zeiss LSM 880 confocal microscope. Z-stack images were analyzed via custom Python scripts that use the Scikit-image library. Three-dimensional images and movies of phage penetration were reconstructed from the Z-stack images via a Python library Mayavi^56^. All data and custom scripts are freely available at https://github.com/krNichith/Phage_Delivery.git

### Fluorescence *in situ* hybridization

5 μL of a day culture of curli fiber producing *E. coli* WR3110 with plasmid pBT1 mCherry was incubated for biofilm production on TSY agar pad for 48 hours at 28°C. 50 μL of *C. gingivalis* suspension was spotted around the circumference of the *E. coli* biofilm and incubated for 24 hours in conditions optimum for *C. gingivalis* motility (described above). After incubation, a thin section of the top layer of agar was sliced and placed on a silane-coated glass slide (Electron Microscopy Sciences, Hartford, PA, catalog # 63411-01) with three GeneFrame chambers (Thermo Scientific, Pittsburgh, PA, catalog # AB0577) stacked on top of the other. 250 μL of PBS (pH 7.1) with 4% of fixing agent paraformaldehyde was poured on the biofilm and the slide was placed in an empty petri dish. A humidity chamber was created by adding 25 mL water to a glass Tupperware box (6” x 4” x 2”). The petri dish containing the slide was incubated for 2 hours at 4°C in the humidity chamber. After incubation, the biofilm was gently washed with 250 μL of PBS. Subsequently, 250 μL of 1 mg/mL lysozyme in 20 mM Tris-HCl (pH 7.5) was added to the GeneFrame chamber and incubated at 37°C for 30 min in the humidity chamber. The lysozyme solution was replaced with a 250 μL hybridization buffer which contained 20 mM Tris-HCl [pH 7.5], 0.9 M NaCl, 20% formamide, 0.01% SDS, and 250 nM of a probe fluorescently labeled with tetrachlorofluorescein (TET). The probe targets the 16S rRNA of *Capnocytophaga* spp. (5’ – TCA GTC TTC CGA CCA TTG – 3’)^31,57^ and was manufactured by Biosearch Technologies, Petaluma, CA. The samples were incubated at 46°C for 4 hours in a modified humidity chamber where 20% formamide was used instead of water. The hybridization solution was removed, and biofilms were washed in 250 μL wash buffer (20 mM Tris-HCl [pH 7.5], 215 mM NaCl, 5 mM EDTA) for 15 min at 48°C. The wash solution was removed, and biofilms were mounted in VectaShield vibrance antifade mounting solution (Fisher Scientific, Pittsburgh, PA, catalog # H-1700-10) for a minimum of 1 hour at 4°C prior to imaging. Z-stack images (3 μm slices) were acquired using a Zeiss LSM 880 confocal microscope and were analyzed using ImageJ. The three-dimensional images and movies of phage penetration were reconstructed via ImageJ.

## Acknowledgements

We thank Lanying Zeng and Knut Drescher for providing strains. This research is supported by NIH R00 grant DE026826.

## Author Contributions

AS and NKR conceptualized the experiments and wrote the paper. NKR performed most experiments and data analysis. AS performed the initial experiment for phage transportation. CAC performed the simulations. AMZ performed the FISH experiment.

## Data Availability

Example datasets and custom data analysis scripts are freely available on GitHub. All additional data are available upon request.

## Supplementary Information

### Supplementary movies

**Movie 1**. Fluorescent lambda phage (green) being actively transported by a swarm of *C. gingivalis* (gray).

**Movie 2**. An example where lambda phage (green) does not adhere to a motile *C. gingivalis* cell.

**Movie 3**. An example where a lambda phage (green) transiently attaches to *C. gingivalis* cell and is transported as cargo.

**Movie 4**. Fluorescent lambda phage (green) diffusing in the thin layer of liquid on the surface of wet agar incubated with 100% relative humidity.

**Movie 5**. Fluorescent lambda phage (green) in the swarm fluid of a *C. gingivalis* with inhibited motility.

**Movie 6**. Time-lapse images of a fluorescent *E. coli* colony (red) in the diffusion control where phages are diffusing in a thin liquid layer on a wet agar surface.

**Movie 7**. Time-lapse image of a fluorescent E. *coli* colony (red) in the experimental setting where phages are delivered by C. *gingivalis* swarms.

**Movie 8**. Time-lapse image of a fluorescent *E. coli* colony (red) in the control where only *C. gingivalis* (no phages) invades the *E. coli* colony.

**Movie 9**. Three-dimensional view of a 425 μm X 425 μm X 105 μm region of curli fiber producing 72 h old *E. coli* biofilm. In this control, phages diffuse in the biofilm and localization of phages after 10 h of diffusion is shown in green.

**Movie 10**. Three-dimensional view of a 425 μm X 425 μm X 105 μm region of curli fiber producing 72 h old *E. coli* biofilm. In this experimental setting, phages are actively transported by *C. gingivalis* and localization of phages after 10 h is shown in green.

**Movie 11**. Three-dimensional view of a curli fiber containing *E. coli* biofilm (red) containing tunnels formed by a *C. gingivalis* swarm (green).

**Movie 12**. Three-dimensional view of the *C. gingivalis (*green) channel from **Movie 11**.

## Supplementary Figure legends

**Fig S1:**
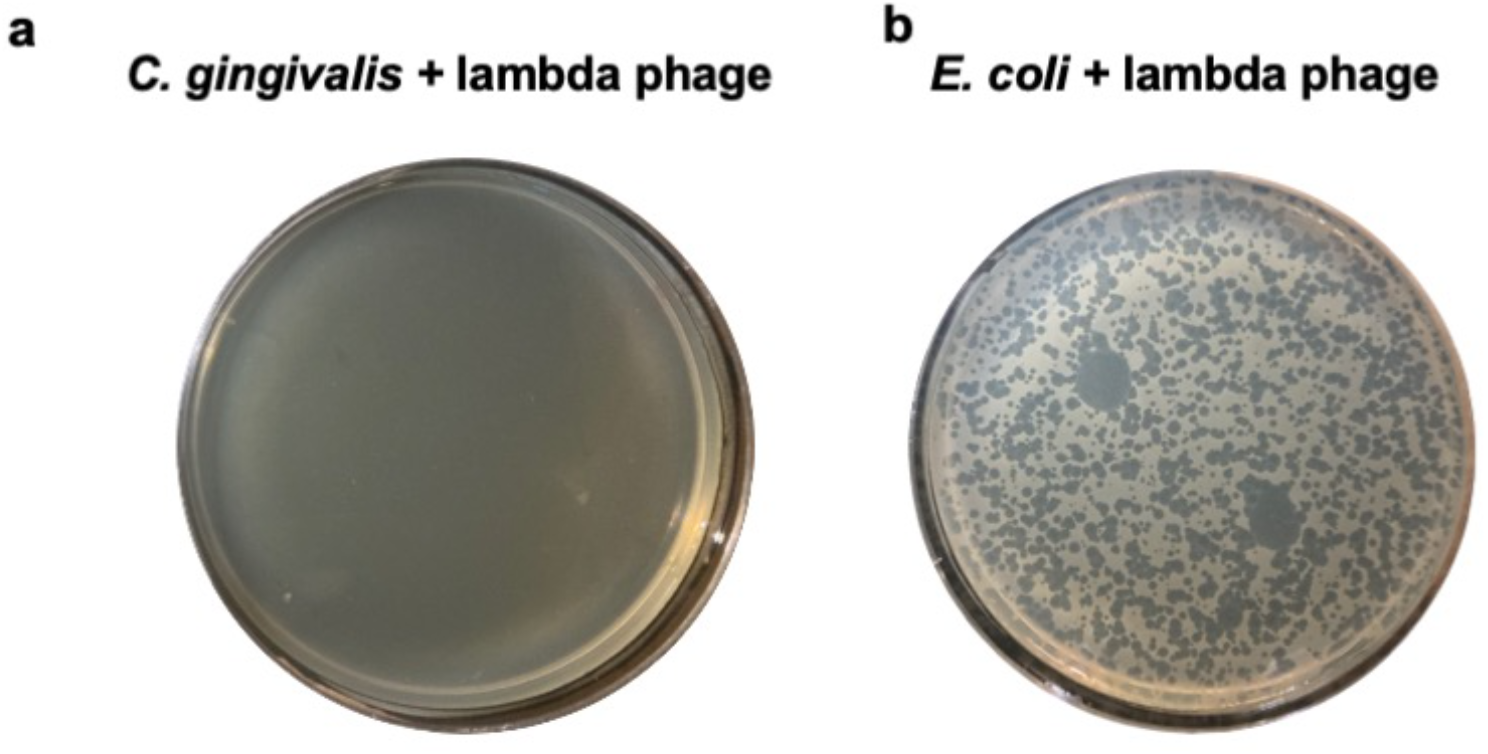
**a**, A plaque assay shows that λ_LZ641_ does not infect *C. gingivalis*. **b**, As a control, a plaque assay shows infection of *E. coli* by λ_LZ641_.

**Fig S2.**
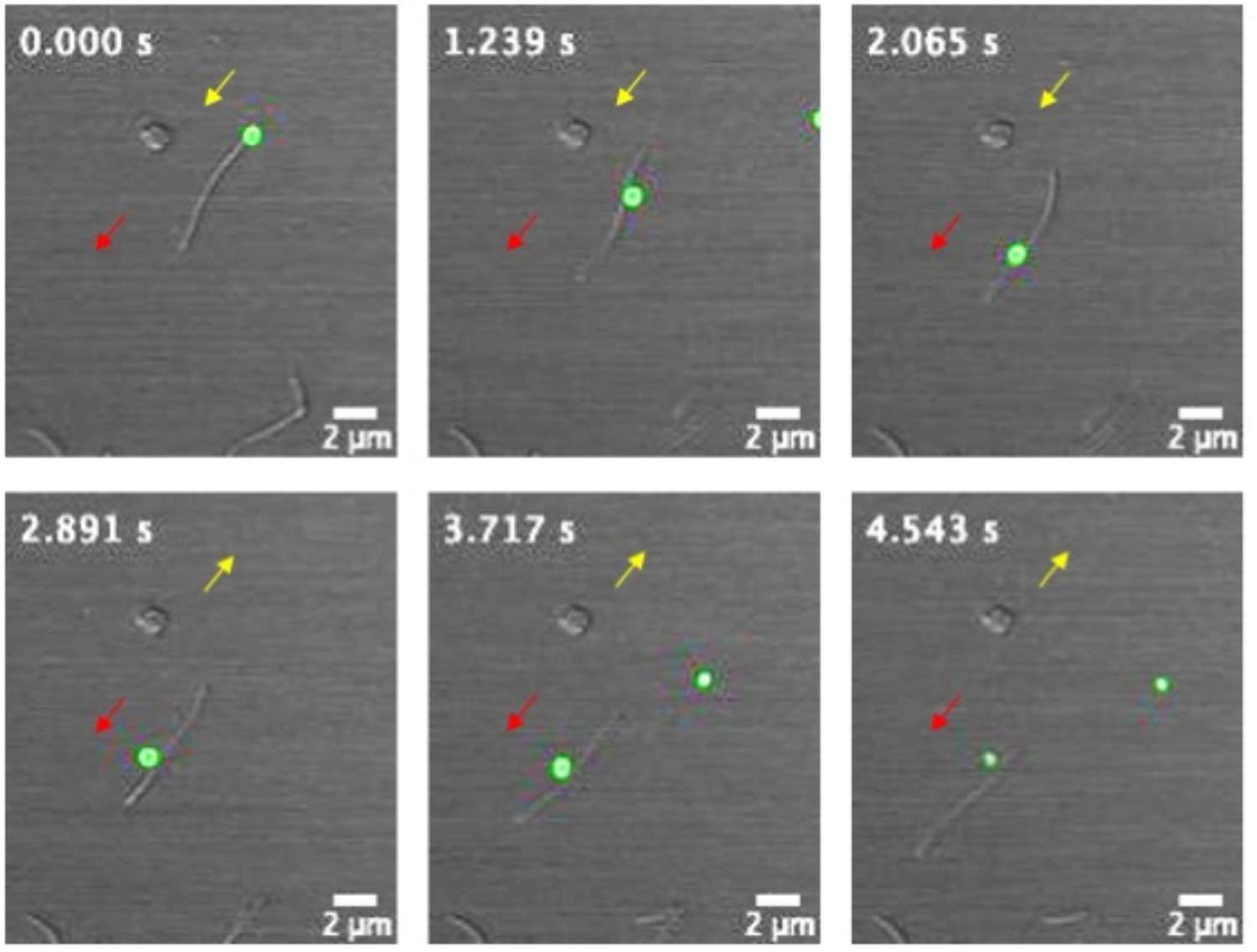
A time-lapse image of a phage particle being transiently propelled along the surface of a *C. gingivalis* cell. Yellow arrow indicates the direction of motion of phage and the red arrow indicates the direction of motion of a *C. gingivalis* cell. The pattern for cell-surface motion of phage is similar to previously observed motion of the cell-surface adhesin SprB.

**Fig S3.**
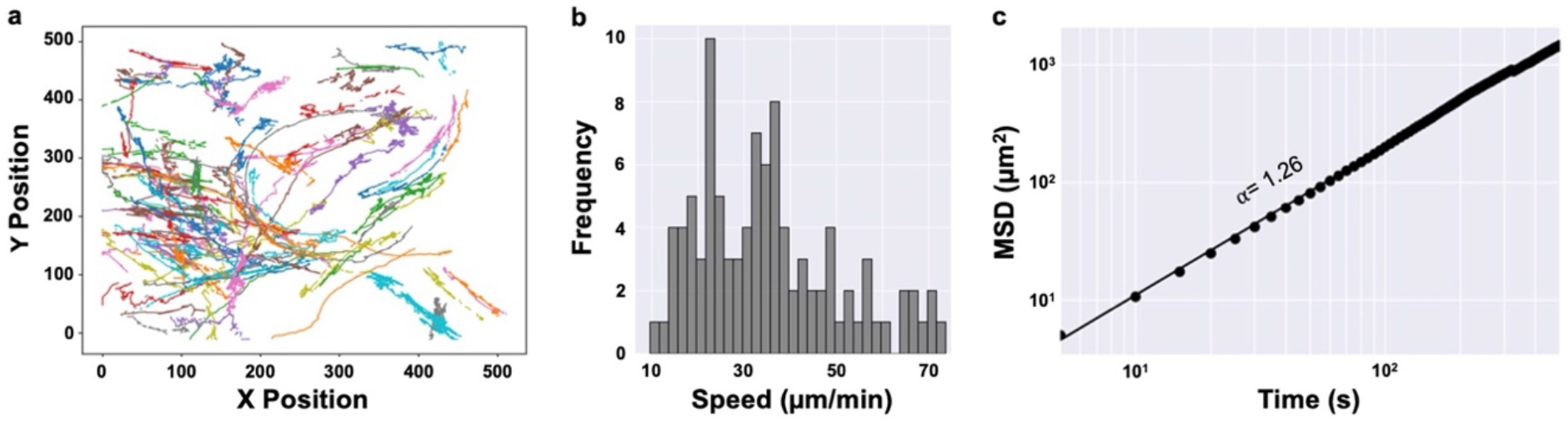
**a**, Combined trajectories from 12 movies with 98 phages propelled by different *C. gingivalis* swarms. **b**, A frequency distribution of the speed at which the 98 phages are transported by *C. gingivalis*. As described previously^21^, the speed is proportional to the number of layers within a swarm. The number of layers within the 12 swarms varies. Hence, we see multiple peaks. **c**, Ensemble mean squared displacement of the 98 phages plotted as a function of time. The slope (α = 1.26) of the power-law fit implies that the phages are super diffusive and are actively propelled by a *C. gingivalis* swarm.

**Fig S4.**
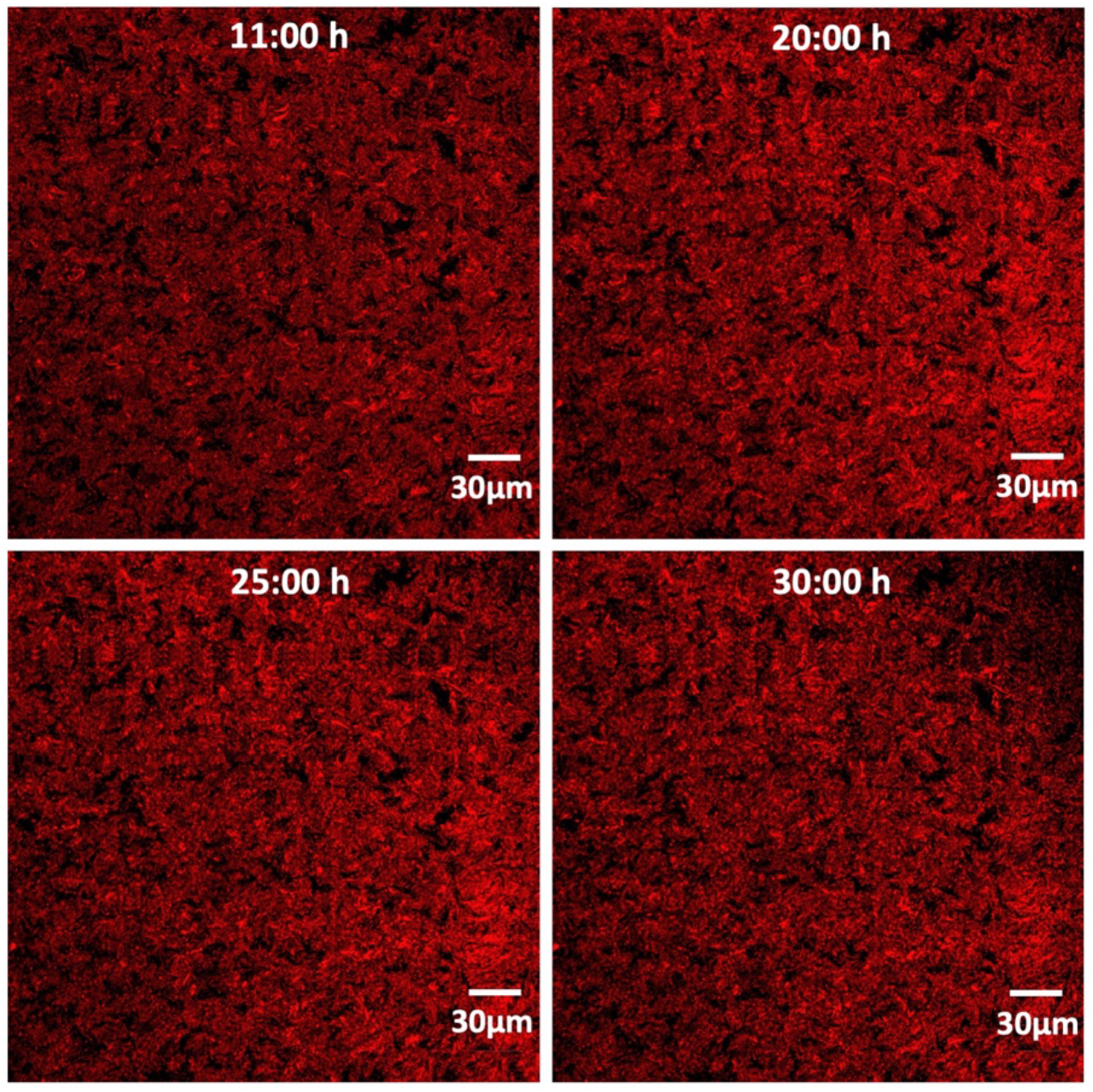
Images of an *E. coli* colony shown at different time points. Here, phages were spotted 1500 μm away from the imaged spot and were diffusing within a thin liquid layer of wet agar (**Movie 6**).

**Fig S5.**
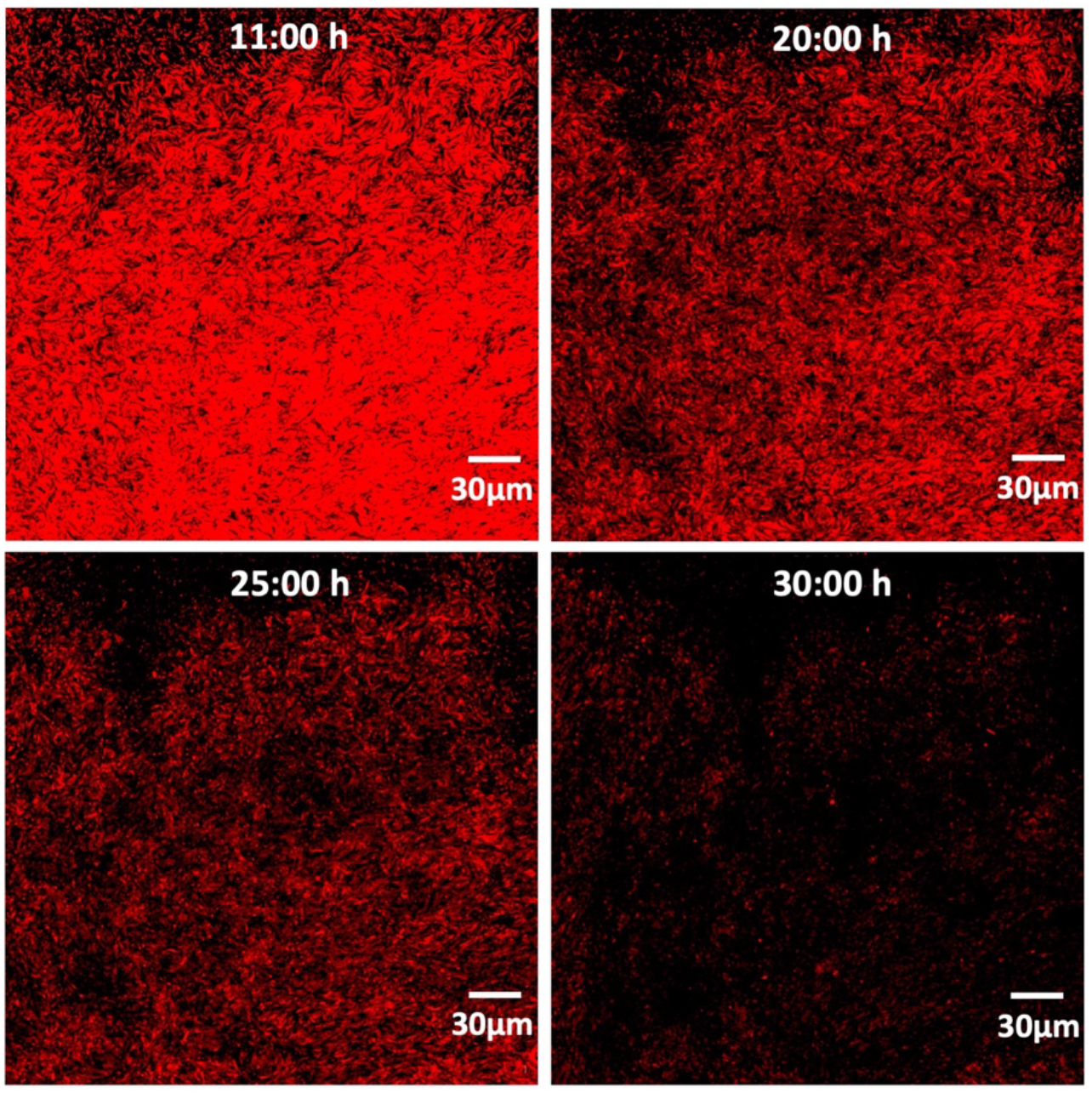
Images of an *E. coli* colony shown at different time points. Here, phage-CG mix was spotted 1500 μm away from the imaged spot and phages were actively delivered by a swarm of C. *gingivalis* (**Movie 7**).

**Fig S6.**
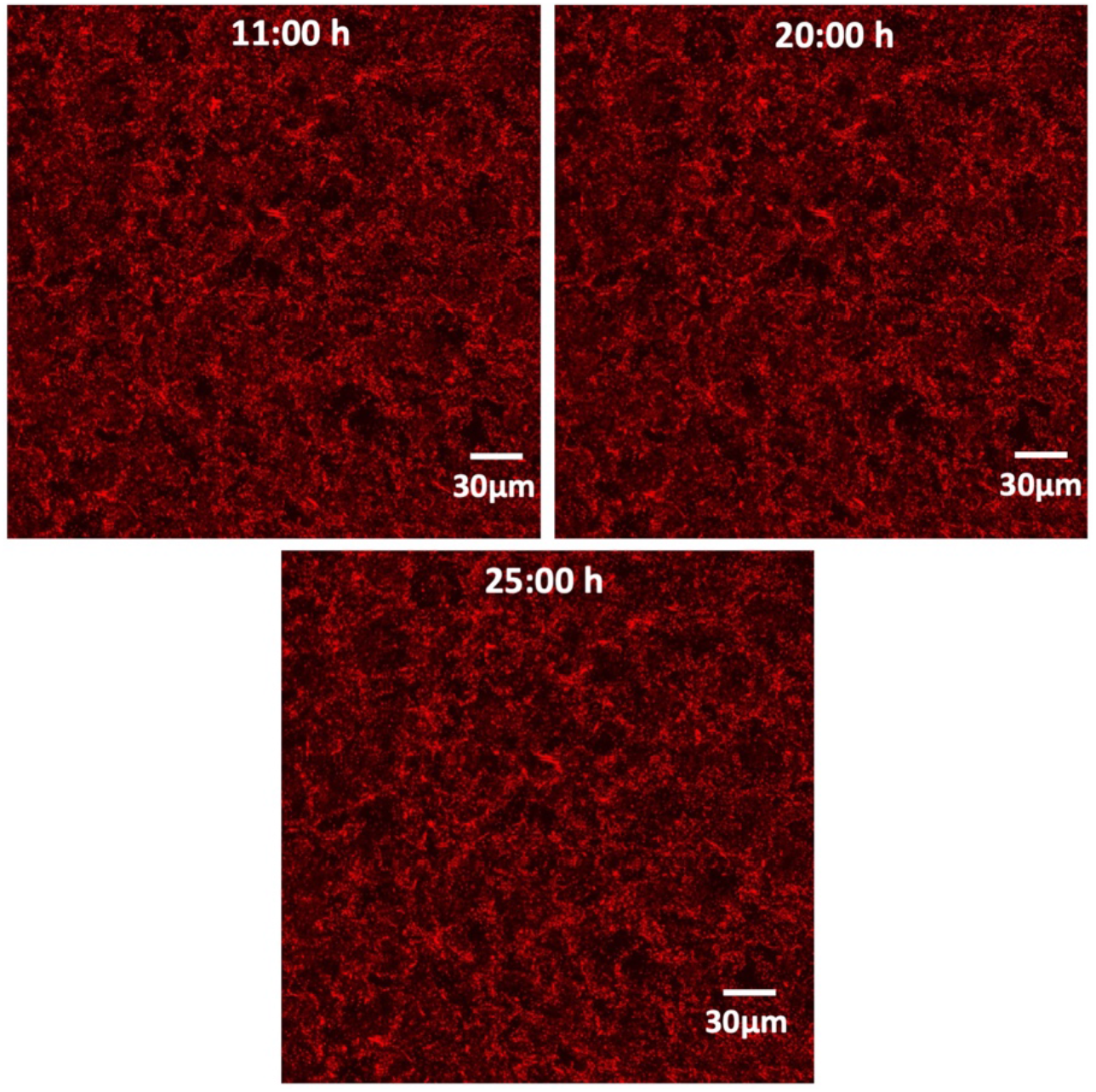
Images of an *E. coli* colony shown at different time points. Here, only *C. gingivalis* cells were spotted 1500 μm away from the imaged spot. No phages were present (**Movie 8**).

**Fig S7.**
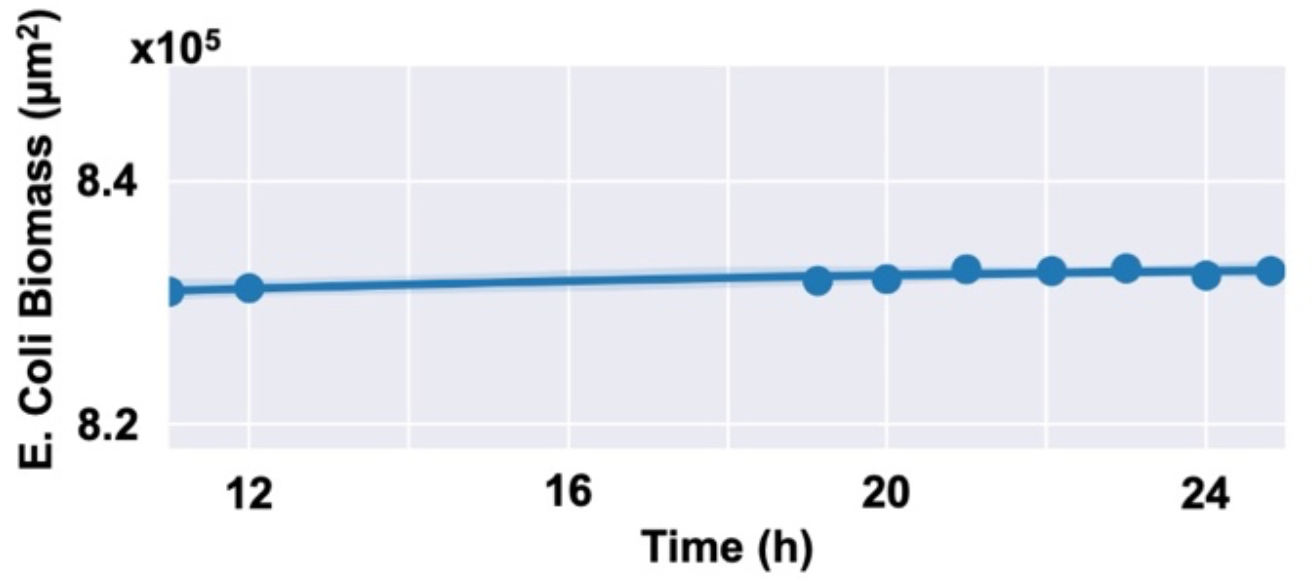
Changes in the area of the *E. coli* biomass depicted as a function of time when only *C. gingivalis* invades an *E. coli* colony (same scenario as **Fig S6**). These measurements are from **Movie 8**. This control shows that there is no change in *E. coli* biomass due to the presence of only *C. gingivalis*.

**Fig S8.**
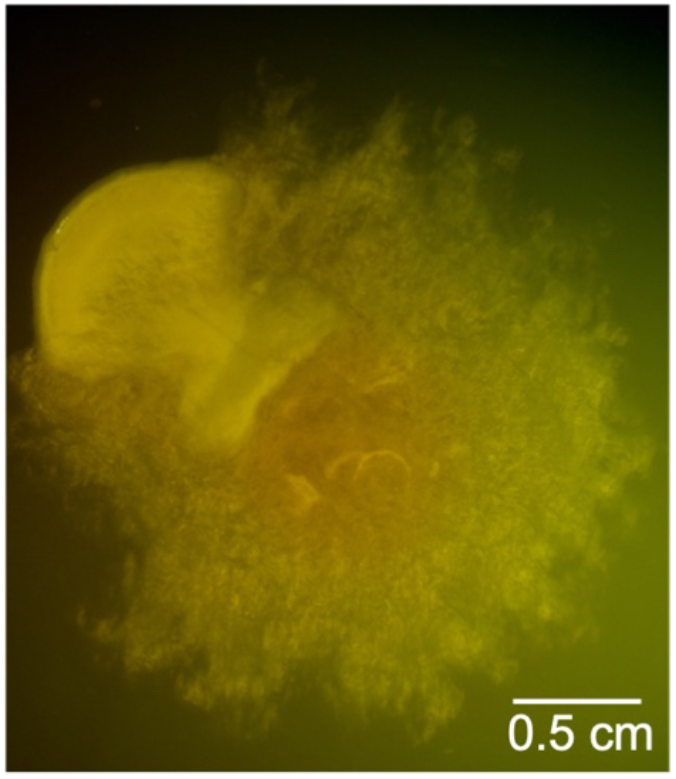
A colony image taken at 30 hours after inoculation showing that *C. gingivalis* (rough) swarms over *E. coli* (smooth).

**Fig S9:**
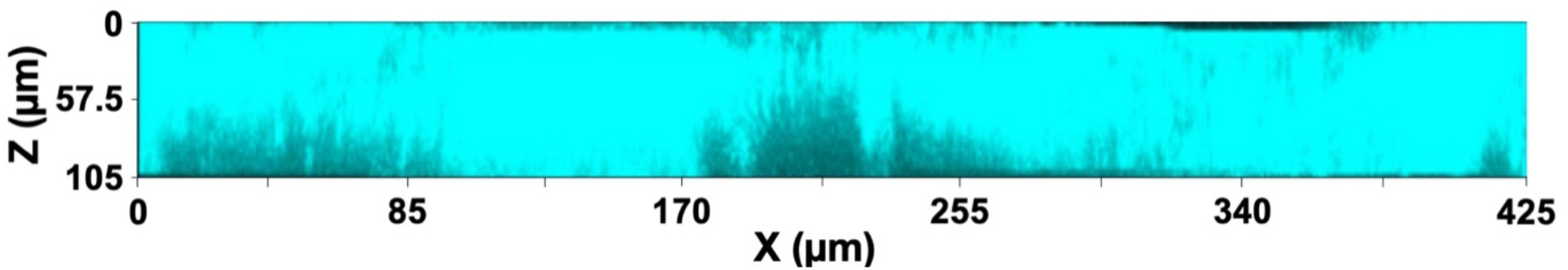
A maximum intensity projection (cyan) of stained curli fiber from z-stack images of a 72 h old *E. coli* biofilm.

**Fig S10.**
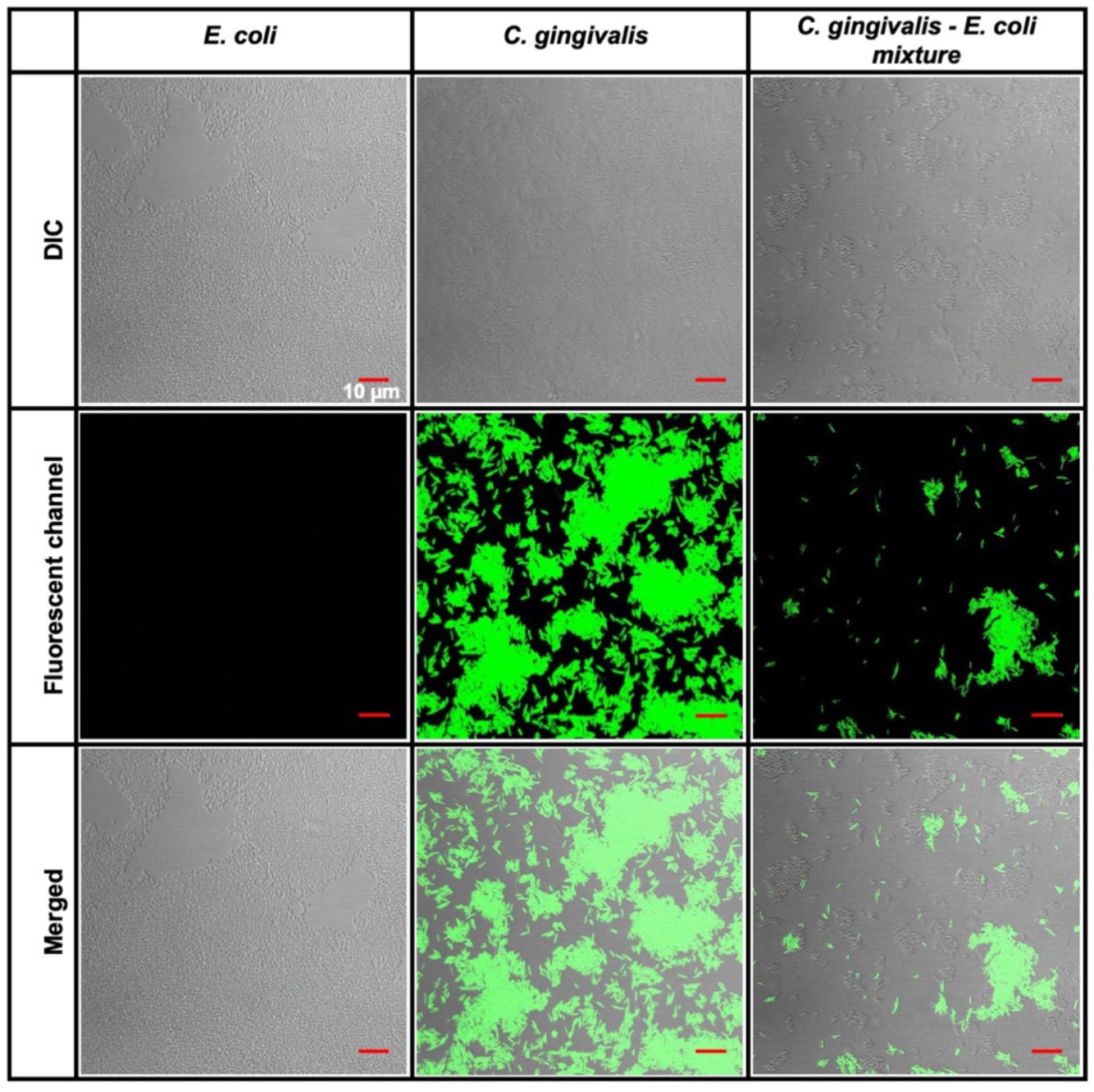
Fluorescent in-situ hybridization (FISH) of cells treated with a probe that targets the 16S rRNA of *Capnocytophaga* spp. Individual images of the DIC and fluorescent channels along with the merged view of the two channels are shown.

## Notes

### Competing Interest Statement

The authors have declared no competing interest.

